# Adhesive interactions within microbial consortia can be differentiated at the single-cell level through expansion microscopy

**DOI:** 10.1101/2024.06.25.600639

**Authors:** Pu-Ting Dong, Wenyuan Shi, Xuesong He, Gary G. Borisy

**Author notes:** **Classification:** Biological Sciences, Microbiology.

## Abstract

**Significance:** A single-cell understanding of microbe-microbe interactions is critical for unraveling the organization and dynamics of microbial communities. Through an unconventional application of expansion microscopy, we oppose the adhesive force holding microbes together by an expansion force pulling them apart, resulting in microbial separation dependent on the strength of microbial adhesion. Our new approach establishes a proof-of-principle for differentiating adhesive interactions within microbial consortia at the single-cell level.

**Abstract:** Investigating microbe-microbe interactions at the single-cell level is critical to unraveling the ecology and dynamics of microbial communities. In many situations, microbes assemble themselves into densely packed multi-species biofilms. The density and complexity pose acute difficulties for visualizing individual cells and analyzing their interactions. Here, we address this problem through an unconventional application of expansion microscopy, which allows for the ‘decrowding’ of individual bacterial cells within a multispecies community. Expansion microscopy generally has been carried out under isotropic expansion conditions and used as a resolution-enhancing method. In our variation of expansion microscopy, we carry out expansion under heterotropic conditions; that is, we expand the space between bacterial cells but not the space within individual cells. The separation of individual bacterial cells from each other reflects the competition between the expansion force pulling them apart and the adhesion force holding them together. We employed heterotropic expansion microscopy to study the relative strength of adhesion in model biofilm communities. These included mono and dual-species *Streptococcus* biofilms, and a three-species synthetic community (*Fusobacterium nucleatum*, *Streptococcus mutans*, and *Streptococcus sanguinis*) under conditions that facilitated interspecies coaggregation. Using adhesion mutants, we investigated the interplay between *F. nucleatum* outer membrane protein RadD and different *Streptococcus* species. We also examined the *Schaalia-TM7* epibiont association. Quantitative proximity analysis was used to evaluate the separation of individual microbial members. Our study demonstrates that heterotropic expansion microscopy can ‘decrowd’ dense biofilm communities, improve visualization of individual bacterial members, and enable analysis of microbe-microbe adhesive interactions at the single-cell level.

## Introduction

Microbes in nature are frequently found as multispecies communities growing on surfaces as biofilms (1–4). The importance of spatial organization for understanding microbial ecosystems (5–8) has been highlighted in recent reviews on the gut microbiome (9, 10), oral microbiome (11–15) and polymicrobial infection (16, 17). Because many microbe-microbe interactions are short range, it is critical to understand spatial organization at the single-cell level. A key approach in meeting this objective has been simultaneous imaging of different bacterial taxa in a complex community through multiplexed fluorescence imaging (18–20). The information obtained from such studies establishes the spatial proximity relationship of individual microbes to other microbes or to host tissue. However, connecting this spatial information to function remains challenging.

One aspect of function is the adhesion of microbes to each other or to host tissue. Adhesion is of fundamental importance in microbial biofilms subject to flow, such as the oral microbiome (21). Because of salivary flow, any oral microbe that does not adhere to host oral tissue or teeth either directly or indirectly via interacting with other adhered microbes will be flushed into the gastrointestinal tract. Adhesion is likely also instrumental to the building of oral biofilms (22) and the emergence of spatial organization as it dictates the stability of microbe-microbe associations. Coaggregation has been used with pure cultures in pairwise combinations *in vitro* to determine the relative tendency of bacterial species to adhere to each other (2, 23, 24). However, this tool while useful as an *in vitro* assay is not suited to evaluating adhesive interactions in a complex, multispecies community *in situ*. Alternative approaches are needed to fill this gap.

Expansion microscopy is a novel approach introduced as a resolution-enhancing method (25). It is based on coupling target molecules or structures to a swellable, polyelectrolyte gel. Isotropic swelling of the gel causes the decrowding of target biomolecules and thus increased resolution with conventional microscopes. The approach has been used to achieve super-resolution imaging of protein complexes (26), RNA complexes (27) and microtubule nanostructures (28). Of note, expansion microscopy also causes the samples to become nearly completely transparent with the refractive index of the expanded gel close to that of water, enabling deep tissue imaging (29).

In this paper, we describe an unconventional application of expansion microscopy to differentiate microbial adhesive interactions at the single-cell level. Whereas in conventional expansion microscopy the objective is to render the gel-embedded tissue as uniform as possible to achieve isotropic expansion, our goal is to maintain the integrity of the microbial cells while expanding the extracellular space between them. The key novel principle we introduce is heterotropic expansion--the application of expansion force to individual cells by embedding them into a swellable gel without swelling the cells themselves. The expansion process, driven by electrostatic repulsion of negatively charged groups on the gel polymer backbone, generates a pulling force on objects contained within the gel, termed as expansion force. Our rationale is that non-interacting or weakly interacting microbes will be pulled apart as manifested by an increase in their inter-microbial distance, whereas strongly adherent cells will resist the expansion force and remain close together (Figure 1). We first evaluated heterotropic expansion using mono-and dual-species *in vitro* biofilms. After establishing minimally perturbing expansion conditions, we applied our approach to investigate the physical interactions in a three-species consortium model consisting of *Fusobacterium nucleatum* (*F. nucleatum*) and two *Streptococcus* species, *Streptococcus mutans* (*S. mutans*) and *Streptococcus sanguinis* (*S. sanguinis*) under conditions that facilitate interspecies co-aggregation. We also investigated an epibiont association, a recently described episymbiosis between an ultrasmall bacterium *Nanosynbacter lyticus* strain TM7x, and its host bacterium, *Schaalia odontolytica* strain XH001 (30). In combination with spectral imaging, our new approach establishes a proof-of-principle for differentiating adhesive interactions within microbial consortia at the single-cell level.

**Figure 1.**
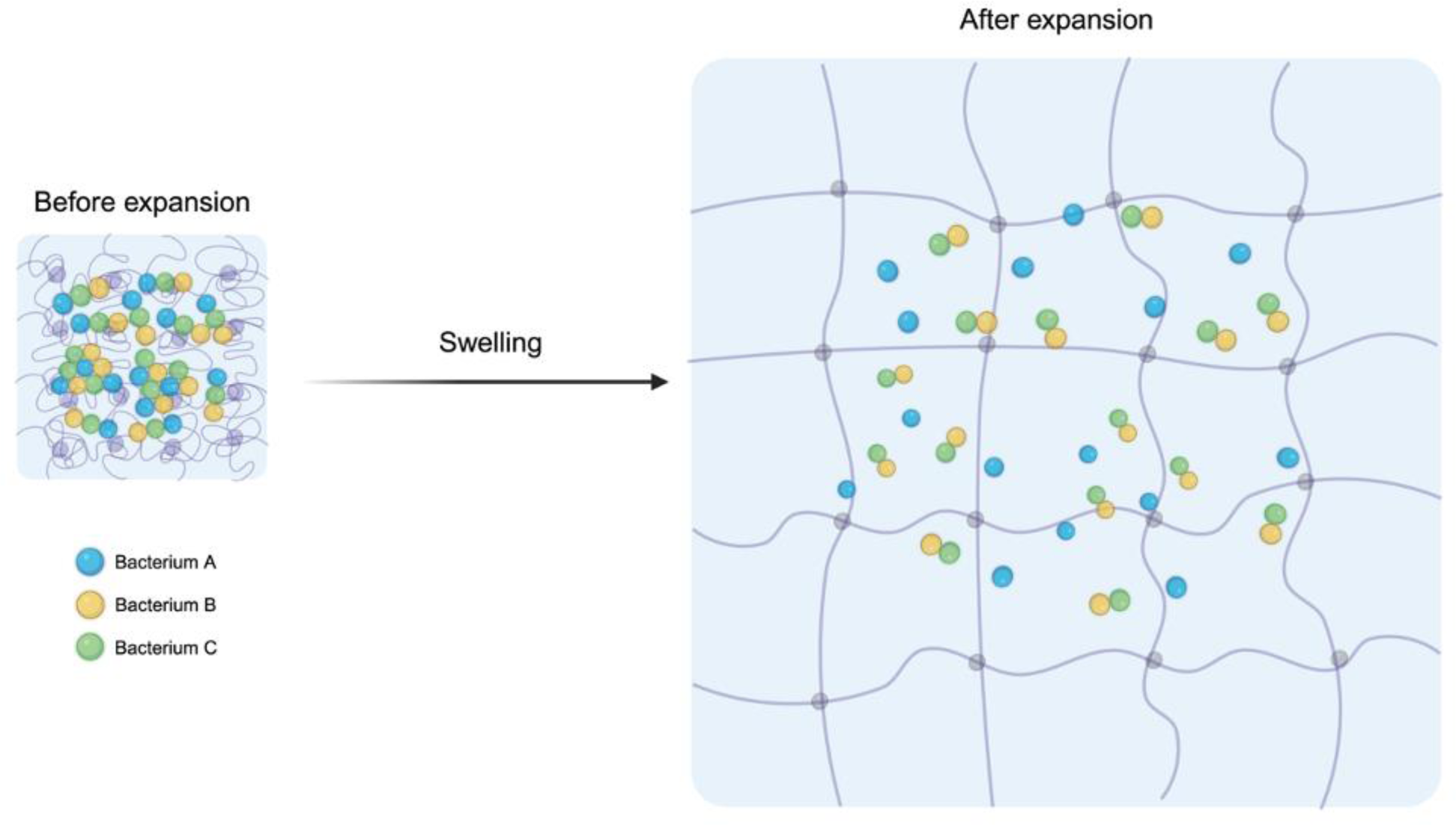
Diagram illustrates application of expansion microscopy for the differentiation of microbe-microbe adhesive interactions within a microbial community. Strongly adhesive bacteria (green and yellow) are not pulled apart during expansion while weakly adhesive bacteria (blue vs green and yellow) are pulled apart.

## Results

### Evaluation of the basic methodology in a mono-species biofilm

Initial experiments were carried out to evaluate whether heterotropic expansion microscopy could be used to decrowd microbial biofilms. This would lay the groundwork for experiments to differentiate microbial adhesive strength within the microbial communities. As a test system, we used *S. mutans* which is known to self-aggregate (31), resulting in biofilms that are spatially heterogeneous. A key property that needed to be established was the behavior of non-interacting objects as a negative control for microbial interactions. For this purpose, fluorescent beads, 2 µm in diameter, were used as a proxy for non-interacting objects. Beads were mixed with *S. mutans* and became incorporated into the mono-species biofilm as the culture grew on saliva-coated glass substrate. The bead-containing biofilm was then embedded in a swellable gel and subjected to the expansion procedure as described in Material & Methods and reported by Chen *et al* (25). In the absence of enzymes such as mutanolysin that specifically digest the bacterial cell wall (32), expansion microscopy can be carried out on microbial cells without compromising cell integrity. During expansion of the biofilm, the distance between beads increased and their concentration per unit area decreased (Figure 2) as expected for a 3-fold expansion of the gel. This result is consistent with the lack of any significant adhesive interaction between the beads.

**Figure 2.**
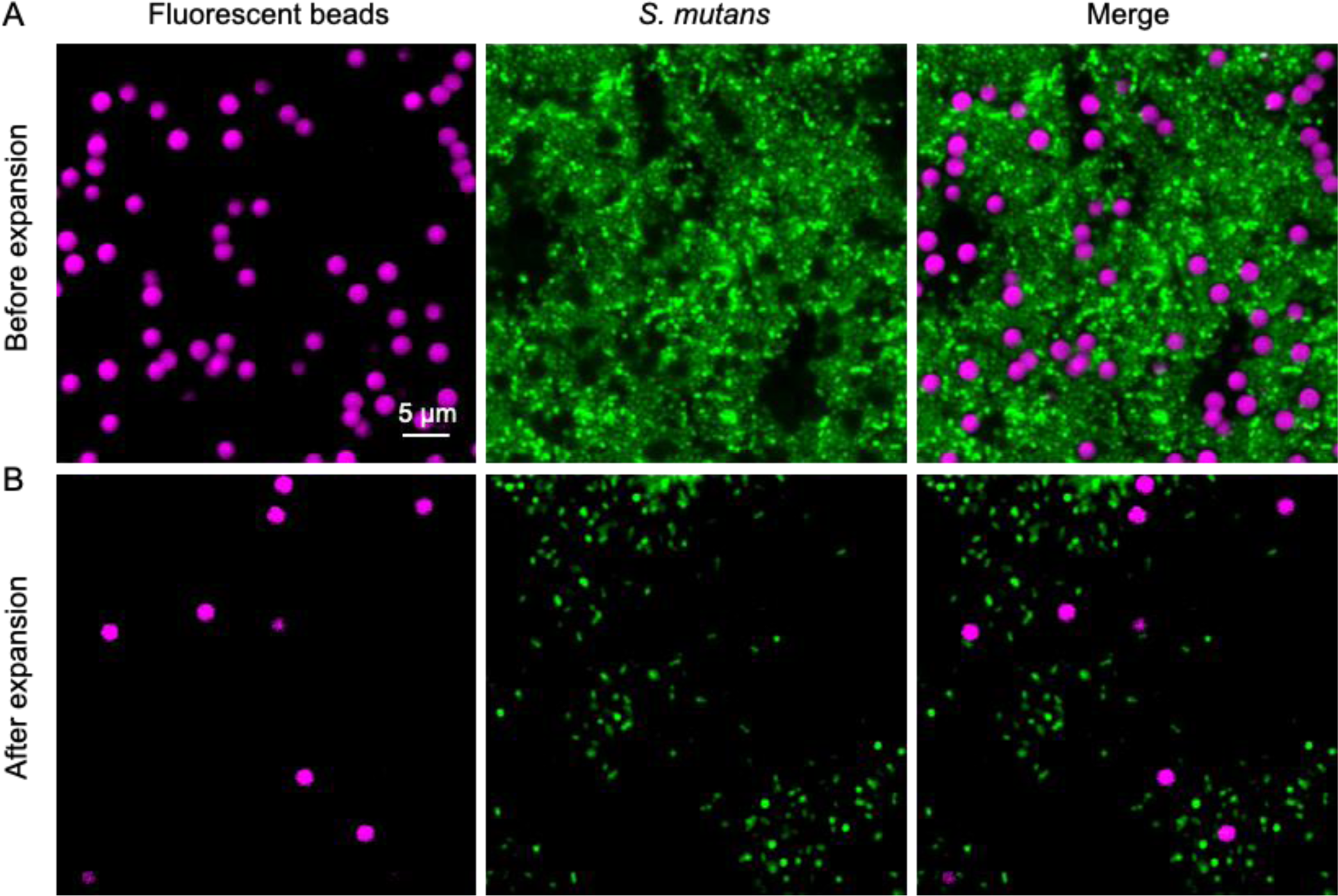
Expansion procedure separates bacteria away from each other in a mono-species biofilm. A representative image of mono-species biofilm before expansion (A) and after expansion (B). Mono-species *S. mutans* (SYTO 9-labeled; green) biofilms were cultured with fluorescent beads (magenta) serving as an internal standard for non-adhesive objects. Images obtained with 63× oil immersion objective (NA=1.4).

Before expansion, the close packing of *S. mutans* in the biofilm hindered visualization of individual cells (Figure 2A). After expansion, *S. mutans* cells showed separation reflecting their initial heterogeneous spatial distribution (Figure 2B). Aggregates of cells evident before expansion were still evident after expansion but were less densely packed as would be expected if the cells were weakly adherent. Importantly, cells were pulled away from each other sufficiently to allow for single-cell visualization. The integrity of the *S. mutans* cells was maintained in the expansion procedure as judged by retention of their size and shape. The separation of individual biofilm cells while maintaining microbial cell integrity establishes that expansion microscopy can be used to decrowd bacterial biofilms.

### Expansion microscopy analysis of a dual-species biofilm

The next question we addressed was whether adhesive interactions between microbes in biofilms could be evaluated quantitatively. For this purpose, we evaluated the application of expansion microscopy to a dual-species biofilm system. When more than one species is present in a biofilm, reporter molecules are necessary to distinguish the different taxa comprising the biofilm. We used genetically encoded fluorescent proteins as reporter molecules for the dual-species biofilm experiments. Biofilms were grown from mCherry-expressing *S. mutans* and GFP-expressing *S. sanguinis* that were mixed 1:1 at an initial OD_600_ of 0.05 in Brain Heart Infusion (BHI) broth. Fluorescent blue beads were added to the mixture as an internal control for measuring the local expansion factor as described before. The fluorescence development of GFP and mCherry is known to require their correct protein folding which is dependent on oxygen (33). Therefore, dual-species biofilms were grown by incubating them aerobically at 37°C for 24 hours.

Imaging of the dual-species biofilms was carried out using spectral imaging followed by linear unmixing to separate the individual fluorescent signals (Materials and Methods). Although not strictly necessary for the dual-species biofilm, we employed the spectral approach in anticipation of subsequent experiments involving multispecies biofilms. Since the growth of *S. mutans* biofilms is known to be strongly dependent on sucrose concentration, we explored growth conditions to achieve a balanced biofilm of both *S. mutans* and *S. sanguinis*. At 1% (w/v) sucrose concentration, *S. mutans* overtook the co-culture, while 0.1% (w/v) sucrose allowed the co-existence of both *Streptococcus* species. Thus, 0.1% (w/v) sucrose was used to grow the dual-species biofilm for expansion analysis.

Using the same expansion procedure as used for the mono-species biofilm, dual-species biofilms were imaged before and after expansion. Before expansion, the biofilm of *S. mutans* and *S. sanguinis* contained closely packed aggregates. Although some individual cells could be discerned, many cells were too close to or overlapped neighbors to allow for unambiguous visualization (Figure 3A). However, after expansion, both *S. mutans* and *S. sanguinis* could be visualized as individual cells in single confocal image planes (Figure 3B), indicating that they were pulled away from each other in the xy plane. Confocal imaging at different axial planes showed that *S. mutans* was more abundant near the glass substrate while *S. sanguinis* was more abundant toward the outer periphery of the biofilm away from the glass bottom of the culture dish. Thus, cells were also pulled apart in the z direction, revealing different spatial distribution patterns for these two streptococci even within the same biofilm.

**Figure 3.**
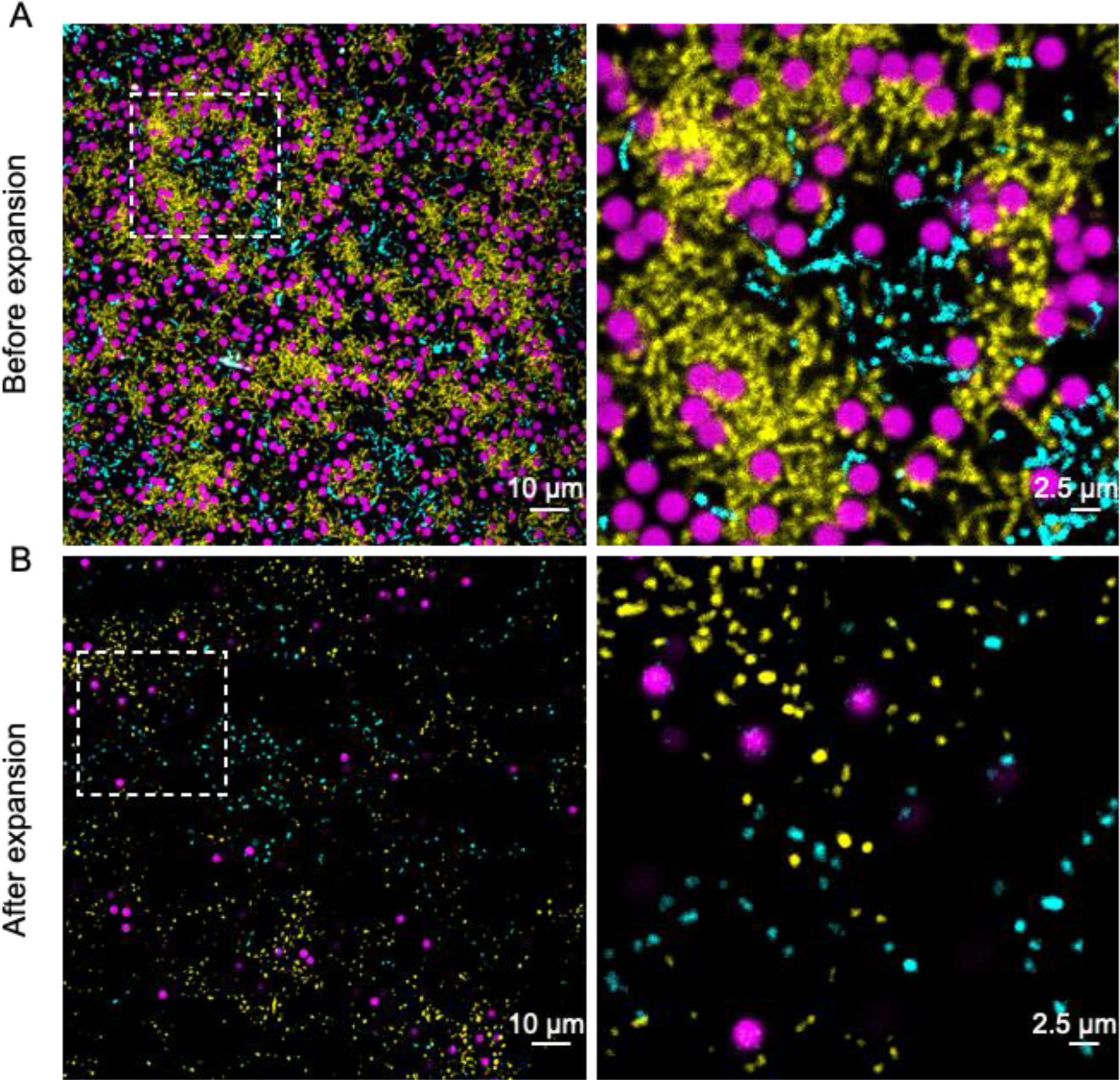
Expansion procedure allows visualization of individual bacteria within a dual-species biofilm. A representative image along with a high-magnification image callout of dual-species biofilm before (A) and after expansion (B). Dual-member *Streptococcus* biofilms were cultured with fluorescent beads serving as an internal standard. Color annotation: fluorescent beads (magenta), mCherry-encoding *Streptococcus mutans* (yellow), GFP-encoding *Streptococcus sanguinis* (cyan).

Spatial proximity relationships of cells within the biofilm were quantified using a program developed for image analysis in microbial ecology, daime (34). Daime quantifies the spatial relationship of two populations by computing a pairwise correlation value as a function of the distance between individual objects. A correlation value larger than 1 means two populations tend to cluster together, less than 1 means they tend to repel each other, and equal to 1 means they are randomly distributed. As shown in Figure 4A, the spatial arrangement of fluorescent beads before and after expansion showed apparent repulsion at short distances and random distribution above a threshold distance. The apparent repulsion can be explained by the physical size of the beads which was 2 µm. Before expansion, the apparent repulsion below 2.5 µm is a consequence of two beads not being able to occupy the same space. After expansion, the threshold of 7.5 µm is a consequence of the linear expansion of approximately 3-fold as found before in the mono-species *S. mutans* biofilm experiments. The random distribution of the beads above these thresholds indicates that they had no intrinsic attraction to or repulsion from each other. Thus, the quantitative analysis of the bead distribution confirmed that the beads may considered as non-interacting, reference objects for interpreting the impact of expansion on the microbes of the biofilm.

**Figure 4.**
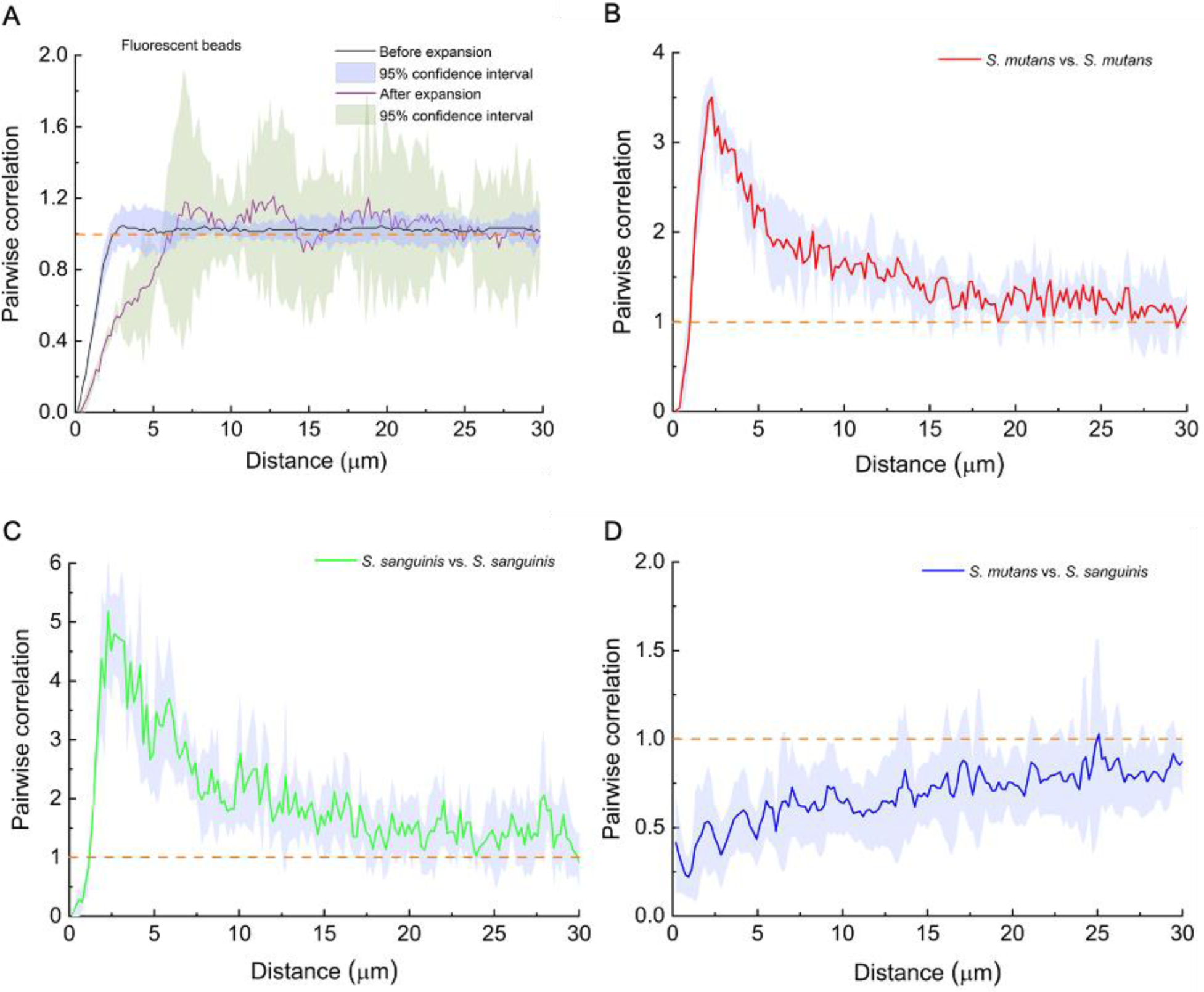
Linear dipole analysis quantifies spatial organization. Proximity analysis of fluorescent beads before and after expansion (A). Linear dipole analysis of images in Figure 3 shows within-taxon autocorrelation (B-C); inter-taxon correlation between *S. mutans* and *S. sanguinis* (D). Data: Mean (solid line) with 95% confidence interval (shaded area). A pair correlation value of 1 is highlighted by orange dashed lines in the figure panels. Pairwise correlations were calculated using 500,000 random dipoles per field of view.

Proximity analysis of microbes in the dual-species biofilm before expansion was ambiguous because many cells in the aggregates overlapped each other and could not be discerned as individual cells. However, after expansion, proximity analysis became unambiguous because individual cells could be clearly visualized. The pairwise correlation patterns of *S. mutans* with itself (Figure 4B) and *S. sanguinis* with itself (Figure 4C) both showed apparent repulsion below 1 µm and positive clustering at greater distances with a peak at about 2.5 µm. Apparent repulsion at short distance is a consequence of the cells, like the beads, not being able to occupy the same space. Positive association at greater distance reflects the existence of close aggregates before expansion which is consistent with the known properties of *S. mutans* and *S. sanguinis* to self-aggregate within biofilms (35). Since expansion was 3-fold and the diameters of *S. mutans* and *S. sanguinis* are about 0.8-0.9 µm, the expected peak in the pairwise correlation curves after expansion was approximately 2.4-2.7 µm, which agrees with the observed values of 2.5-3.0 µm. This result indicates that many *S. mutans* and *S. sanguinis* cells were in close association before expansion and were pulled apart by the expansion process. The pairwise correlation between *S. mutans* and *S. sanguinis* was less than 1 over most of the distance range, suggesting apparent repulsion (Figure 4D). However, this negative association does not indicate an intrinsic repulsive interaction between these two *Streptococcus* species; rather, it is a mathematical consequence of the positive clustering behavior of each of the *Streptococcus* species taken individually.

The dual-species biofilm confirmed important components of our expansion methodology. These included the compatibility with genetically encoded fluorescent proteins as reporters, the feasibility of imaging expanded gels by spectral imaging, and the quantification of spatial proximity relationships by pairwise correlation analysis. The results quantitatively demonstrate that microbial associations in a biofilm can be investigated at the single-cell level by expansion microscopy.

### Differentiation of microbial interactions in a three-species consortium model

The results with mono-and dual-species biofilms provided the groundwork for applying heterotropic expansion microscopy to analyze adhesive interactions within multispecies microbial communities. As a model system, we adopted an *in vitro* three-member (*F. nucleatum*-*S. mutans*-*S. sanguinis*) consortium under conditions that facilitate interspecies coaggregation. The binding of different bacteria to each other via specific adhesion molecules (36, 37) can be visually assayed by the resulting formation of flocculant precipitates (38). Coaggregation is thought to play an essential role in the development of multispecies communities, such as dental plaque biofilms (39, 40). However, coaggregation is a population-level phenotype which is not a direct measurement of the adhesive interactions between individual bacteria. Therefore, it would be significant to deconstruct the coaggregation phenotype at the single-cell level.

To differentiate adhesive interactions within a multispecies bacterial system, we designed a minimally perturbing expansion protocol. All three bacterial species were labeled vitally. Wild type *F. nucleatum* was labeled with a membrane-specific fluorophore, FM 1-43. The labeled *F. nucleatum* was mixed with mCherry-expressing *S. mutans* and GFP-expressing *S. sanguinis* at a ratio of 1:1:1 in coaggregation buffer (Materials and Methods). The mixture resulted in the formation of flocculant precipitate which settled at the bottom of the tube. Samples of the precipitated aggregates were transferred to a glass bottom dish, and gelation was conducted directly on the aggregates. This minimally perturbing protocol differed from conventional expansion protocols in three ways: no fixation was carried out; no Acryloyl-X molecular linker was employed; and no proteinase K digestion was carried out before expansion. Thus, in this protocol, the samples were essentially unfixed, unlinked and undigested. Images before expansion were taken immediately after the gelation step. After gelation, expansion was driven by adding water to the aggregates-containing gel and changing the water three times over a period of 1-2 hours. The size of the aggregates increased after expansion but were less densely packed as would be expected from the separation induced by the expansion process.

Before expansion (Figure 5A), even under a high-magnification 63× oil immersion objective (NA=1.4) with a pixel size of 65 nm, we could not discern single bacteria inside the aggregates, let alone determine the three-dimensional architecture of these aggregates. However, after expansion, even with a low-magnitude 20× objective (NA=0.8), individual bacteria were clearly distinguishable in single-plane confocal images (Figure 5B). We also examined the distribution of bacteria within the aggregates in the axial direction by acquiring images along the z-axis and representing the results through orthoslice views (Figure 5C). The results show that all three bacterial species could be resolved individually in both lateral and axial directions.

**Figure 5.**
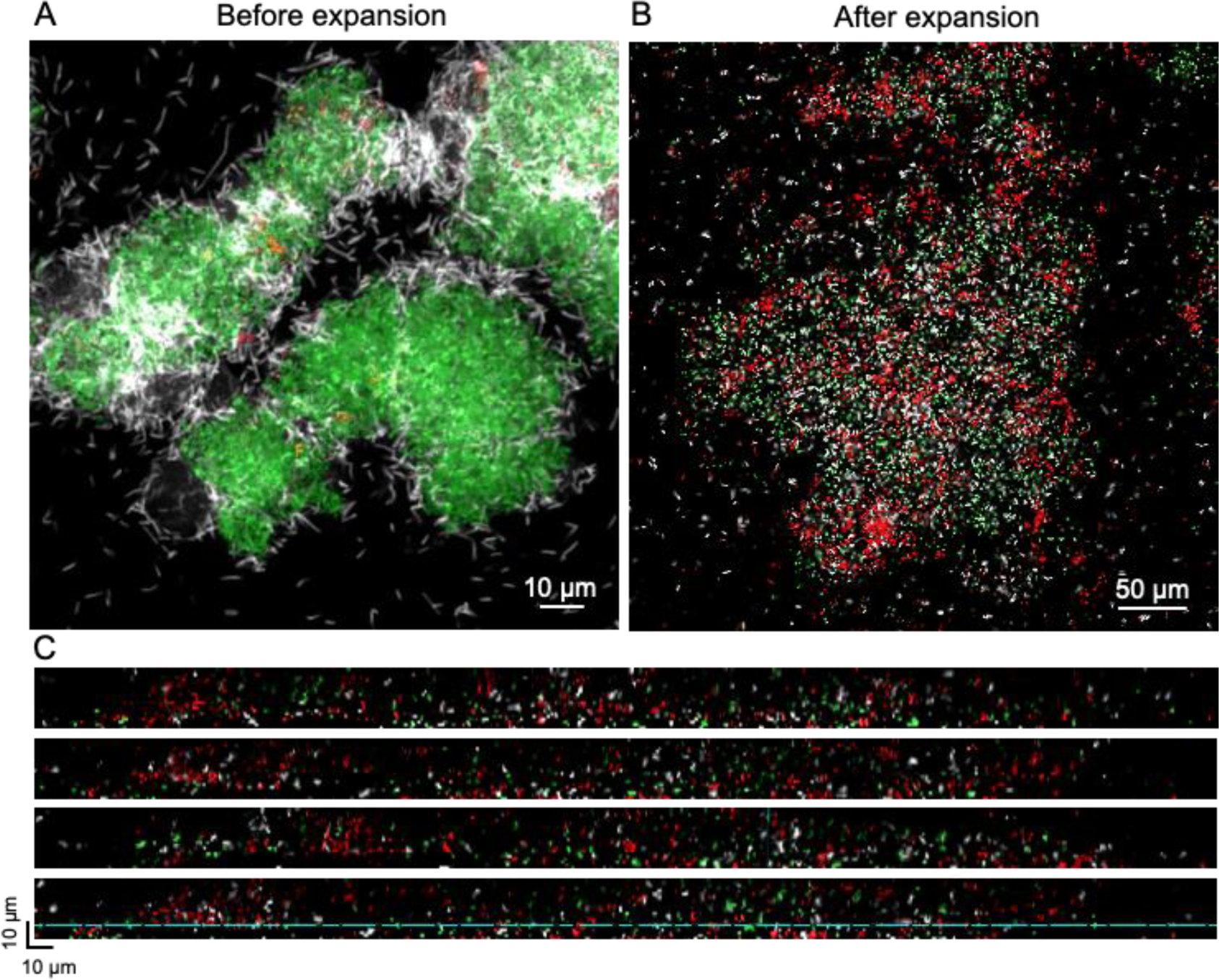
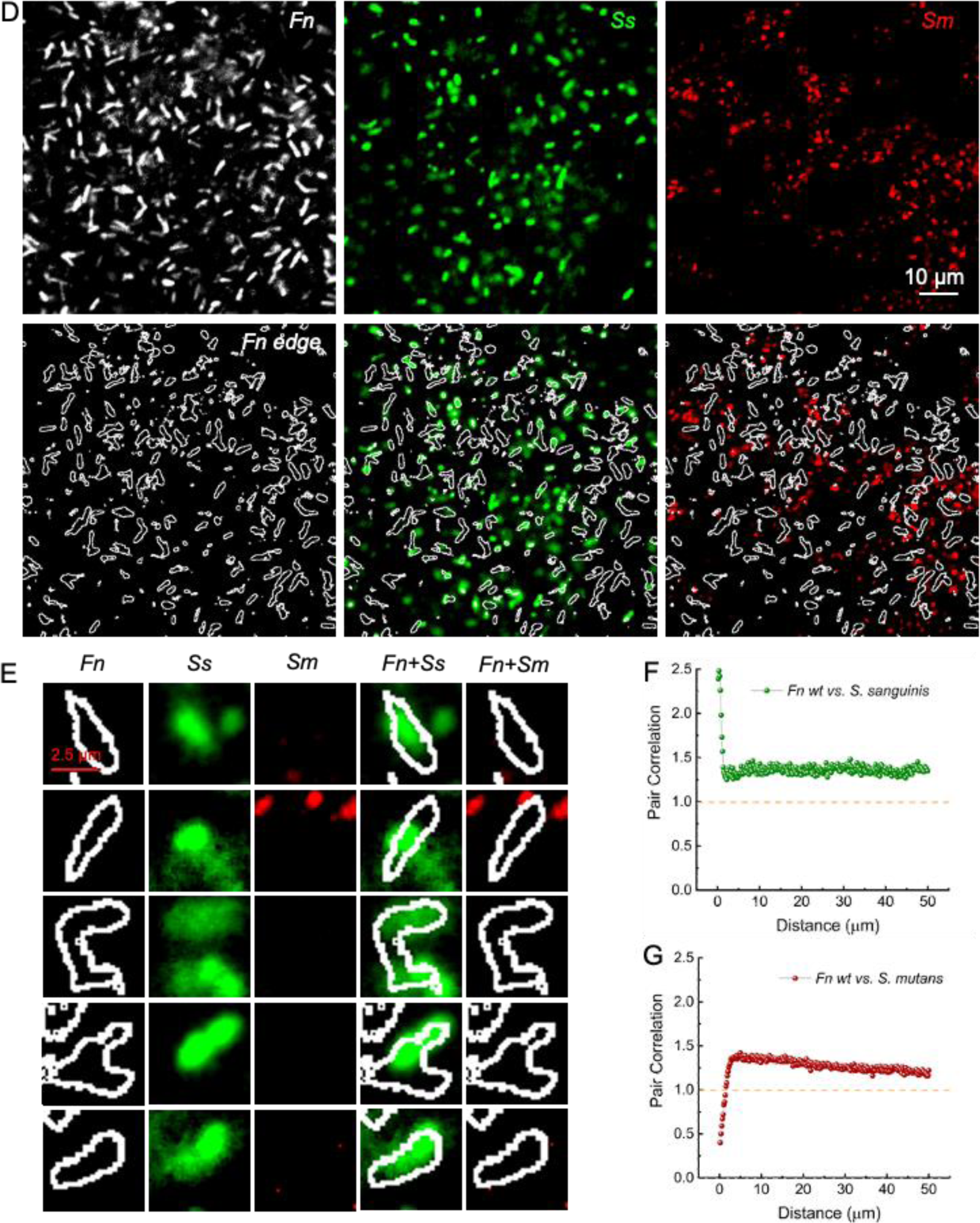
Expansion microscopy differentiates the interaction between *F. nucleatum* wildtype with two *Streptococcus* species. (A). Spectral imaging (after linear unmixing) of *F. nucleatum-S. mutans-S. sanguinis* aggregates before expansion under a 63× oil immersion objective (NA=1.4). Scale bar = 10 µm. (B). Spectral imaging (after linear unmixing) of *F. nucleatum-S. mutans-S. sanguinis* aggregates after expansion. 20× objective (NA=0.8). Scale bar = 50 µm. (C). Orthoslice views of *F. nucleatum-S. mutans-S. sanguinis* aggregates in panel (B) in the axial and lateral directions. (D-E). High-magnification views (D) and single-cell image callouts (E) of merged view of *F. nucleatum* wildtype with *S. mutans* and *S. sanguinis*. (F-G). Proximity analysis of image in (D) between *F. nucleatum* and different *Streptococcus* species aggregates after expansion. A pair correlation value of 1 is highlighted by orange dashed lines in the figure panels. Pairwise correlations were calculated using 500,000 random dipoles per field of view. Color annotation: *F. nucleatum* (white), *Streptococcus mutans* (red), *Streptococcus sanguinis* (green).

The capacity to resolve individual cells through expansion microscopy enabled us to perform quantitative proximity analysis at the single-cell level on the model consortium. Proximity analysis demonstrated differential interactions between *F. nucleatum* and *S. sanguinis* vs *F. nucleatum* and *S. mutans. F. nucleatum* showed a strong association with *S. sanguinis* at very short distances (< 3 µm) but essentially a random association at greater distances (Figure 5D, F). In contrast, *F. nucleatum* showed a repulsion to *S. mutans* at very short distances, changing to a neutral interaction at greater distance (Figure 5D, G). The remarkable difference in behavior of *F. nucleatum* with respect to *S. sanguinis* and *S. mutans* was investigated further by acquiring high-magnification views of the aggregates after expansion. As shown in Figure 5D, the close association of *F. nucleatum* and *S. sanguinis* was frequently manifested as overlapping cells. Overlapping *S. sanguinis* and *F. nucleatum* can be documented in two ways: first, from direct visualization of high magnification views, individual and merged panels show that *F. nucleatum* is juxtaposed with *S. sanguinis* (Figure 5E); second, from the proximity analysis (Figure 5F-G), the high positive pairwise correlation value at short distances (< 0.5 µm) indicates that the bacterial cells are closer together than a single cell diameter. In contrast, *F. nucleatum* and *S. mutans* were nearby or adjacent to each other but still clearly distinct and generally not overlapping. The exclusion at short distances accounts for the apparent repulsion of *F. nucleatum* and *S. mutans* shown by the proximity analysis. We interpret these images and the proximity analysis results to signify that *F. nucleatum* was in tight association with *S. sanguinis* but not with *S. mutans.* In short, our heterotropic expansion protocol indeed captured differences in the adhesion behavior of different bacterial species in our model system at the single-cell level.

### Evaluation of *F. nucleatum* RadD dependent and independent adhesion to different *Streptococcus* species

The ability of *F. nucleatum* to adhere to other bacterial members of dental plaque has been attributed to a group of outer membrane adhesion proteins (41). Among these, RadD has been identified as the primary mediator of the inter-species adherence between *F. nucleatum* and various *Streptococcus spp*., including *S. mutans* and *S. sanguinis* (42, 43). Here, we evaluate the adhesion of wild type *F. nucleatum* and *F. nucleatum* Δ*radD* which is depleted in RadD using heterotropic expansion microscopy.

The three-species consortium containing *F. nucleatum* Δ*radD*, *S. mutans* and *S. sanguinis* aggregated less in comparison to consortia with *F. nucleatum* wildtype (Figure 6A), confirming a role for RadD in coaggregation. Nonetheless, some aggregation occurred. The residual aggregation with the *F. nucleatum* Δ*radD* suggests a RadD-independent mode of coaggregation as well. This result is in accord with the literature that adhesin(s) in addition to RadD may also contribute to *F. nucleatum*-*Streptococcus* coaggregation (41). We then analyzed whether microbe-microbe interaction inside these non-RadD aggregates could be differentiated from those in the wild type consortia. Individual cells were clearly distinguished after expansion inside these aggregates (Figure 6B), which was further confirmed by images of orthogonal views. High magnification merged images of *F. nucleatum* Δ*radD* and the two *Streptococcus* species presented a similar association pattern as that of *F. nucleatum* wildtype: a larger number of *S. sanguinis* cells overlapped spatially with *F. nucleatum* compared to *S. mutans* (Figure 6C-D). Proximity analysis (Figure 6E-F) showed that both *Streptococcus* species, especially *S. sanguinis* demonstrated a close association with *F. nucleatum* Δ*radD* while the relationship between *S. mutans* and *S. sanguinis* appeared spatially random (see SI Fig. S1C).

**Figure 6.**
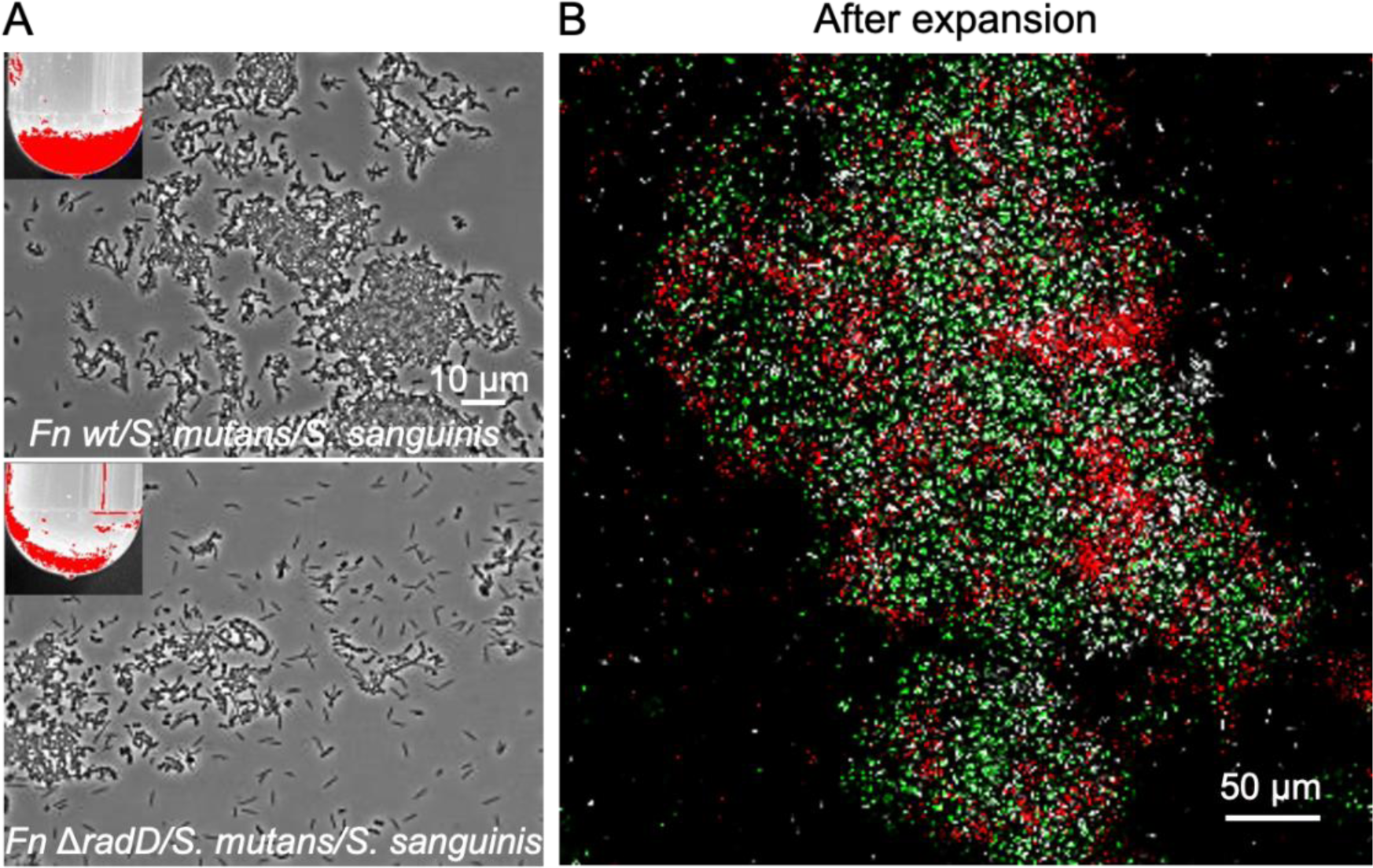
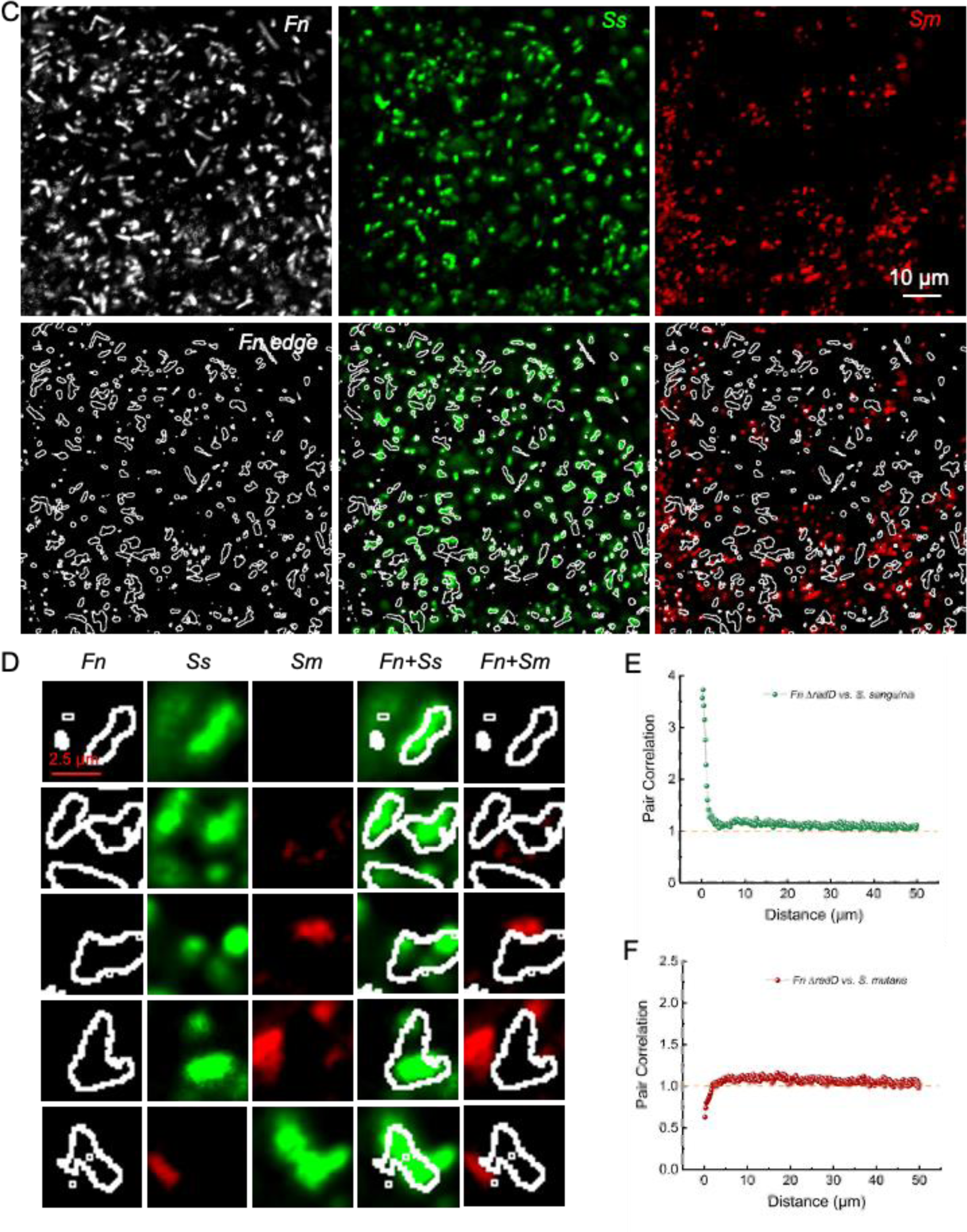
Expansion microscopy differentiates the interaction between *F. nucleatum* Δ*radD* with two *Streptococcus* species. (A). Coaggregation assay comparison between *F. nucleatum* wildtype and *F. nucleatum* Δ*radD* with two different *Streptococcus* species. Images obtained by phase microscopy; insert shows the aggregation in test-tube with heat-map to help visualize flocculant precipitate. (B). Spectral imaging (after linear unmixing) of *F. nucleatum* Δ*radD-S. mutans-S. sanguinis* aggregates after expansion under a 20× objective (NA=0.8). Scale bar = 50 µm. (C-D). High-magnification views (C) and single-cell image callouts (D) of merged view of *F. nucleatum* Δ*radD* with *S. mutans* and *S. sanguinis*. (E-F). Proximity analysis of image in (C) between *F. nucleatum* Δ*radD* and different *Streptococcus* species aggregates after expansion. A pair correlation value of 1 is highlighted by orange dashed lines in the figure panels. Pairwise correlations were calculated using 500,000 random dipoles per field of view. Color annotation: *F. nucleatum* Δ*radD* (white), *Streptococcus mutans* (red), *Streptococcus sanguinis* (green).

To gain a quantitative comparison of the spatial arrangement within the aggregates from the three-species consortium containing *F. nucleatum* wildtype or Δ*radD* in terms of the interaction with *S. mutans* and *S. sanguinis*, we randomly chose eight different fields of view to conduct the proximity analysis and then compared them statistically. As shown in SI Fig. S1A, there was no apparent difference in the pairwise correlation values between *F. nucleatum* wildtype/*S. mutans* and *F. nucleatum* Δ*radD/S. mutans*. Similarly, the pairwise correlation values between *F. nucleatum* Δ*radD* and *S. sanguinis* was almost the same as that of *F. nucleatum* wildtype with *S. sanguinis* (SI Fig. S1B) while the spatial relationship between *S. mutans* and *S. sanguinis* was essentially random (SI Fig. S1C). Each of the three taxa, *F. nucleatum*, *S. mutans* and *S. sanguinis,* showed self-correlation at short distances indicating a tendency to self-associate (SI Fig. S1D-F). Taken together, these data suggest that a non-RadD adhesin or adhesins contribute to the interaction between *F. nucleatum* and *Streptococci* in addition to the RadD adhesin. Furthermore, the non-RadD adhesin-mediated interaction between *F. nucleatum* and *S. sanguinis* was of sufficient strength that it was not disrupted by the expansion force.

To understand whether the repulsive behavior between *S. mutans* and *F. nucleatum* (both wildtype and *radD* mutant) at short distances in three-species consortium is due to the competitive association between *S. sanguinis* and *F. nucleatum*, we carried out coaggregation experiments and expansion analysis between *F. nucleatum* and individual *Streptococcus* species under the same experimental conditions. Proximity analysis (SI Fig. S2) showed that both *F. nucleatum* wildtype and *F. nucleatum* Δ*radD* displayed close association with *S. sanguinis* at short distances. In contrast, the association with *S. mutans* appeared to be repulsive at shorter distances and random at greater distances. Collectively, this evidence suggests that the close association between *F. nucleatum* and *S. sanguinis* is intrinsic, and the apparent repulsive behavior between *F. nucleatum* and *S. mutans* is not from competition with *S. sanguinis*.

### Expansion microscopy analysis of a microbial episymbiosis model

Epibionts, by definition, are organisms that live on the surface of another organism and, therefore, have a close structural association. We reasoned that the association of a bacterial epibiont with its host cell might be particularly strong. As an additional illustration of the application of heterotropic expansion microscopy, we examined a two-member oral episymbiotic system in which the epibiont was ultrasmall Saccharibacteria *Nanosynbacter lyticus* strain TM7x and its host was a *Schaalia odontolyticus* strain XH001 (30, 44).

To query whether the expansion force could separate TM7x away from its host cell XH001, we applied the expansion procedure to cocultures vitally stained with SYTO-9 (Materials and Methods). Low magnification images of the coculture (Figure 7A) show aggregates of cells in which individual XH001-TM7x complexes were hard to discern. After expansion, XH001-TM7x complexes were pulled away from each other but many TM7x cells remained closely attached to their host cells (Figure 7B) suggesting that the interaction force between TM7x and its host cell is stronger than the expansion force. High-magnitude images confirmed that expansion did not detectably alter the physical association between TM7x and its host cell (Figure 7B inserts).

**Figure 7.**
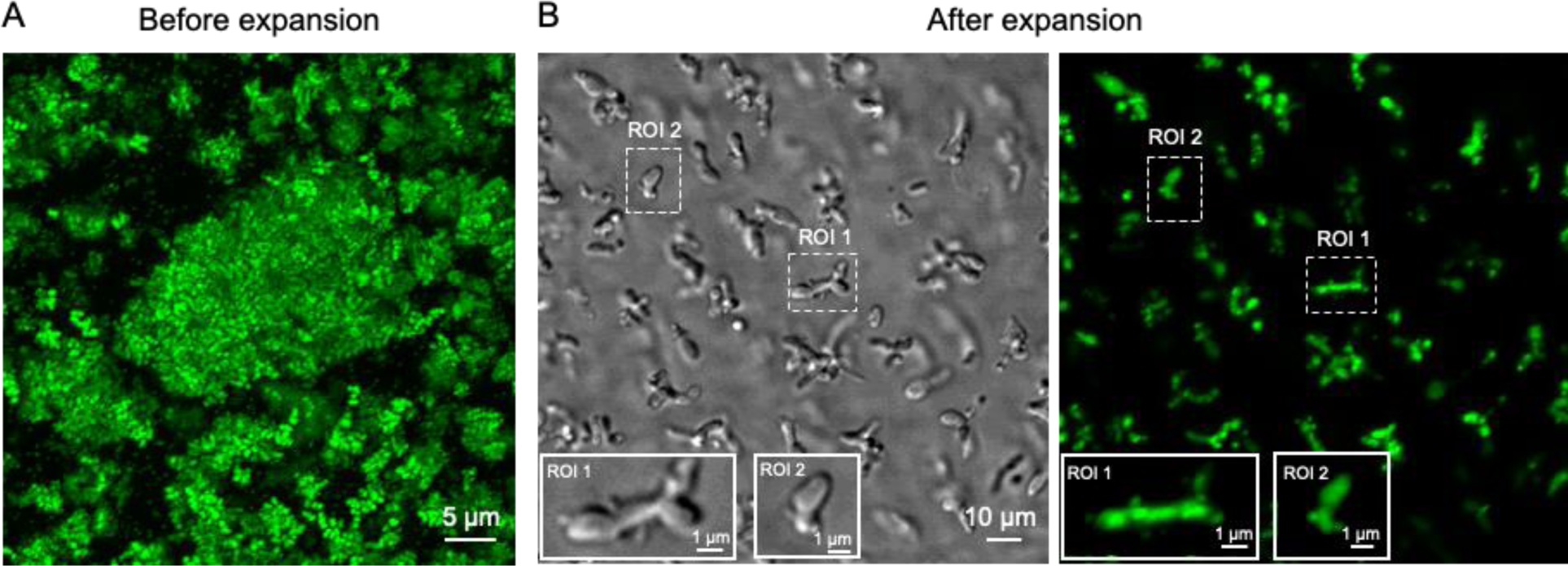
Expansion force does not pull apart the epibiont interaction pair: Saccharibacteria *Nanosynbacter lyticus* strain TM7x and *Schaalia odontolytica* (XH001). Confocal fluorescence imaging of SYTO 9-labeled coculture XH001-TM7x complexes before (A) and after expansion (B). Two regions of interest (ROI), dashed white boxes, are shown at high magnification (inserts).

## Discussion

Our study demonstrates the utility of an unconventional application of expansion microscopy to the analysis of microbial biofilms. We show how heterotropic expansion can ‘decrowd’ dense biofilm communities, improve visualization of individual bacterial members, and enable differentiation of microbe-microbe adhesive interactions within microbial communities at the single-cell level.

The application is unconventional in that its purpose is to exploit the force driving expansion to assess microbial adhesion rather than to achieve super-resolution imaging. In conventional expansion microscopy, the biological matter is digested to enable uniform, isotropic expansion of the polymer gel. In contrast, in our application, the microbes remain intact while being entrapped in the polymer network. The space between the microbes is expanded while the space within the microbes is not, hence the term heterotropic expansion. Expansion of the swellable polymer gel generates a pulling force on the entrapped microbes. The result is a ’tug-of-war’ competition between the expansion force and microbial adhesive forces. Weakly adherent cells will be pulled apart while strongly adherent cells will remain together. Thus, heterotropic expansion enables the differentiation of microbe-microbe interaction within microbial communities at the single-cell level.

Our application of expansion microscopy is complementary to the conventional coaggregation assay, which assesses the binding of pairs of bacteria to each other by whether they form visible flocculant precipitates. Heterotropic expansion microscopy deconstructs the coaggregation phenotype at the single-cell level. Specifically, through investigating the aggregates formed by the model community, *F. nucleatum*-*S. sanguinis*-*S. mutans,* we found that the spatial interaction between *F. nucleatum* and *S. sanguinis* was significantly different from that of *F. nucleatum* and *S. mutans*, a feature otherwise not evident in the coaggregation assays. The *F. nucleatum*-*S. sanguinis* adhesion was evidently stronger than the expansion force since cells remained in close proximity and many were juxtaposed upon each other even after expansion. In contrast, the aggregates formed between *F. nucleatum* and *S. mutans* were pulled apart by expansion, indicating that the *F. nucleatum*-*S. mutans* adhesion was weaker than the expansion force. The epibiont association between TM7 and *Schaalia odontolytica* is another example of a tight association as the expansion process failed to pull apart the interacting partners. This result was not surprising since the association between an epibiont and its host was expected to be strong. Studies have shown that type IV pili trigger a close association between Saccharibacteria and its host bacteria (45).

RadD, a major outer membrane protein of *F. nucleatum,* has been reported to play a key role in the binding phenotype between *F. nucleatum* and *Streptococci* including *S. mutans* (46) and *S. sanguinis* (41) and, consequently, in the integration of *F. nucleatum* into the supragingival microbial community (42). We investigated whether RadD was essential for the strong adhesive interaction between *F. nucleatum* and *S. sanguinis* by testing a RadD deletion mutant of *F. nucleatum.* Although the three-species community, *F. nucleatum* Δ*radD*-*S. sanguinis*-*S. mutans,* showed a reduced level of coaggregation, expansion microscopy revealed that binding between *F. nucleatum* Δ*radD* and *S. sanguinis* remained tight. This result indicated that RadD was not essential for tight binding at the single-cell level and that a non-RadD adhesin or adhesins had sufficient binding force to underlie the tight binding observed. By either mechanism, the strong association between *F. nucleatum* and *S. sanguinis* supports the premise that this interspecies interaction serves as an important association in the organization of supra-and subgingival plaque (47, 48).

Although RadD was not essential for the tight binding between *F. nucleatum* and *S. sanguinis,* deletion of RadD did lead to significant reduction in the overall amount of coaggregation. How can these apparently conflicting results be explained? One possible explanation arises from the likely multivalency of binding interactions. Each binding partner, *F. nucleatum* or *S. sanguinis,* bears multiple copies of adhesin ligands or receptors. The large aggregates and flocculant precipitate resulting from mixture of the two partners may be understood in a manner similar to an antibody-antigen precipitation reaction. Following the antibody-antigen paradigm, we varied the concentration ratio between *F. nucleatum* and *S. sanguinis* in the coaggregation experiment and found that both *F. nucleatum* Δ*radD* and *F. nucleatum* wildtype had similar coaggregation profiles (SI Fig. S3), peaking at a ratio (*F. nucleatum*/*S. sanguinis*) of one. However, the amount of coaggregation with *F. nucleatum* Δ*radD* was approximately half of that for *F. nucleatum* wildtype. Coaggregation at a *F. nucleatum* Δ*radD/S. sanguinis* ratio of 1.0 was essentially the same as that of *F. nucleatum* wildtype at a *F. nucleatum*/*S. sanguinis* ratio of 0.5. From this result, we infer that the reduced coaggregation observed after deletion of RadD may be a consequence of reduction in the number of adhesins per *F. nucleatum* and, therefore, its effective valency ratio to *S. sanguinis.* In sum, the reduced level of aggregates seen by coaggregation is fully consistent with the tight binding between *F. nucleatum* Δ*radD* and *S. sanguinis* as assayed at the single-cell level.

Electro-repulsion among the negatively charged carboxylic groups in the swellable gel generates the expansion force according to Coulomb’s law. However, the net expansion force will become attenuated gradually as it is opposed by the elasticity of the polyelectrolyte backbone, resulting in a balance of force and a stable extent of expansion. It is noteworthy that the initial expansion force is adjustable depending on the concentration of monomer and crosslinker molecules during the gelation step (49). Therefore, we envision that gels with a series of expansion forces can be prepared by carefully choosing the gelation recipes. In this way, we can better differentiate microbial adhesive forces within microbial communities by varying the expansion force.

Our approach for differentiating adhesive interactions within microbial consortia at the single-cell level has some limitations. First, microbial consortia often involve multiple bacterial species, where multiplexed fluorescence imaging is needed. Our study investigated adhesive interactions in a model microbial community consisting of three members. To investigate the feasibility of this single-cell functional platform within complex microbial communities, it will be necessary to evaluate the compatibility between heterotropic expansion microscopy and multiplexed fluorescence imaging. Second, our approach thus far achieved the differentiation of interspecies microbial adhesive strengths on a relative basis. However, the absolute binding strengths between microbes with their neighbors remain to be determined. For the quantification of microbial binding forces, a calibration method will be needed to quantify the gel expansion force. Finally, we need to take into consideration whether the expansion process affects the viability of microbes. Notwithstanding these limitations, our new approach establishes a proof-of-principle for differentiating adhesive interactions within microbial consortia at the single-cell level.

## Materials and Methods

### Materials

Acryloyl-X SE (Invitrogen, A20770), Triton X-100 (Sigma Aldrich, 9036-19-5), DMSO (Sigma Aldrich, 67-68-5), EDTA, disodium (0.5 M, pH=8, Invitrogen, AM9260G), Tris·HCl (1 M, pH=8, ThermoFisher Scientific, 15568025), NaCl (Fisher Scientific, AC447302500), proteinase K (1500 U/ml, ThermoFisher Scientific, 25530049), UltraPure^TM^ TEMED (ThermoFisher Scientific, 15524010), ammonium persulfate (APS, ThermoFisher Scientific, 17874), sodium acrylate (Sigma Aldrich, 408220-25G), calcium chloride (Sigma Aldrich, 10043-52-4), magnesium chloride (Sigma Aldrich, 7786-30-3), sodium azide (Sigma Aldrich, 26628-22-8), glycine (ThermoFisher Scientific, 36435.A1), sucrose (Fisher Scientific, AA3650830), acrylamide (40% (wt/vol), Fisher BioReagents, BP1402-1), N,N’-methylenebisacrylamide (Sigma Aldrich, 110-26-9), Hi-Di^TM^ formamide (Fisher Scientific, 44-407-53), phosphate buffered saline, PBS (10× stock, Fisher BioReagents, BP3991). Poly-L-lysine solution (Sigma Aldrich, 25988-63-0). 2-µm fluorescent beads (fluorescent blue, Sigma Aldrich, L0280; fluorescent red, Sigma Aldrich, L3030). Dextran, Alexa Fluor^TM^ 647 (ThermoFisher Scientific, D22914). SYTO^TM^ 9 (ThermoFisher Scientific, S34854). FM^TM^ 1-43 (ThermoFisher Scientific, T3163). High-grid glass bottom dish (ibidi, 81148). Glass bottom dish (Fisher Scientific, NC0662883).

### Bacteria and culture

#### Mono-species *Streptococcus mutans* biofilms

Planktonic *S. mutans* UA159 was cultured in Brain Heart Infusion (BHI) broth under aerobic (21% O2) conditions. To prepare *S. mutans* biofilms, *S. mutans* (OD_600_=0.05) was mixed with 20 µl of fluorescent blue beads into fresh BHI (supplemented with 0.1% sucrose) and added to saliva-precoated glass bottom dish. Robust biofilms were formed after overnight aerobic incubation.

#### Dual-species *Streptococcus spp.* biofilms

GFP-expressing *Streptococcus sanguinis* (*S. sanguinis*) and mCherry-expressing *S. mutans* were cultured in BHI broth under aerobic (21% O_2_) conditions to allow maturation of GFP and mCherry. Then, *S. mutans* (OD_600_=0.05) and *S. sanguinis* (OD_600_=0.05) were mixed with 20 µl of fluorescent blue beads into fresh BHI (supplemented with 0.1% and 1% sucrose, respectively) and added to saliva-precoated glass bottom dishes. Robust dual-species biofilms were formed after overnight aerobic cultivation.

#### *F. nucleatum-S. mutans-S. sanguinis* coaggregation assay

*Fusobacterium nucleatum* ssp. *nucleatum* strain ATCC 23726 was cultured in Columbia Broth (Fisher Scientific, DF0944-17-0) under anaerobic conditions (5% H_2_, 5% CO_2_, 90% N_2_) at 37℃. Then, *F. nucleatum* 23726 (*F. nucleatum* wildtype) was stained with a membrane-labeling fluorophore, FM^TM^ 1-43 for 30 min. Then, stained *F. nucleatum* were washed with 1×PBS before combining with the two *Streptococcus* species (mCherry-expressing *S. mutans* and GFP-expressing *S. sanguinis*) in the coaggregation buffer (see recipe). Three bacterial species were then mixed together thoroughly at an OD_600_ of 2 for each bacterial species and allowed to stand in the dark. Flocculant precipitates appeared within 30 min after mixing. Precipitated aggregates were removed three hours after gently mixing and placed onto a glass bottom dish, semi air-dried, followed by gelation, imaging before expansion, expansion by solvent exchange with water, and imaging after expansion.

## Solutions

**Coaggregation buffer:** were prepared according to reference (50). NaCl (150 mM), Tris·HCl (1 mM, pH=8.0), CaCl2 (0.1 mM), NaN3 (0.02% (w/v)), MgCl_2_ (0.1 mM) were mixed in sterile water.

**Expansion-related solutions:** were prepared as reported (29).Acryloyl-X SE (**AcX**) stock solution: 5 mg of acryloyl-X SE was dissolved in 500 µl of anhydrous DMSO. Aliquots were stored at -20℃.

**TEMED** and **APS stock** solutions: both chemicals were dissolved in water at a concentration of 10 g/100 ml, and then divided into aliquots of 1 ml each and stored at -20℃.

Monomer solution (**Stock X**): Sodium acrylate (9% w/v), acrylamide (2.7% v/v), N,N’-methylenebisacrylamide (0.16% w/v), NaCl (12.4% w/v), in 1×PBS. The Stock X solution was then divided into aliquots of 940 µl each and stored at -20℃.

**Gelation solution:** Stock X solution plus TEMED (0.2% w/v), APS (0.2% w/v).

**Digestion buffer:** 0.5% w/v Triton X-100, 1 mM EDTA, 50 mM Tris·HCl, NaCl (4.67% w/v), proteinase K (15 U/ml).

Digestion buffer was also divided into aliquots of 1 ml each and stored at -20℃, proteinase K was added to the remaining digestion buffer immediately before the digestion step.

### Gelation, digestion, expansion on biofilms

*S. mutans* UA159 biofilms and dual-species *Streptococcus* biofilms were fixed in 10% formalin (Sigma Aldrich, HT501128-4L) for 1 hour, then glycine (100 mM) was added to quench the fixation for 5 min. After that, fixed biofilm samples were washed with 1×PBS twice before the gelation step. *S. mutans* UA159 biofilms were then stained with SYTO 9. 0.1 mg/ml AcX in PBS (dilute AcX stock into PBS) was immediately added to the fixed biofilms samples overnight. After incubation with AcX, biofilms were washed with PBS twice, each time for 15 min. Then monomer solution (see recipe) was added to biofilms and gelation was allowed to occur for 1 hour at 37℃. Immediately afterwards, digestion was carried out in the presence of proteinase K (10 U/ml, see recipe). After overnight-digestion, expansion was induced by adding water to the digested samples, and exchange with water for three times, each time for 20 min. Part of the expanded samples were then carefully mounted to a poly-L-lysine-coated cover glass (VWR, 16004-348), and a few drops of water were added to the top of the gel before image acquisition to prevent sample dehydration.

### Gelation, expansion of F. nucleatum wildtype/F. nucleatum ΔradD-S. mutans-S. sanguinis aggregates

Gelation in the presence of the monomer solution (Stock X) was immediately applied to the semi-dried three-member aggregates at 37℃ for 1 hour at a humidified chamber. Spectral imaging was then employed after gelation to acquire images before expansion. After imaging, water was directly added to the gels and gel samples were exchanged with water three times, each time for 20 min. Part of expanded samples were then carefully mounted to a poly-L-lysine coated cover glass (VWR, 16004-348), and a few drops of water were added to the top of the gel before image acquisition to prevent sample dehydration over the process of image acquisition.

### Gelation, expansion of epibiont interaction pair Saccharibacteria *Nanosynbacter lyticus* strain TM7x/*Schaalia odontolytica* (XH001)

Preparation of epibiont pair XH001/TM7x was described in the previously published studies (30, 51). Coculture was then harvested by passaging for two times, washed with 1×PBS. Cells were stained with SYTO 9 afterwards, and washed with 1×PBS. SYTO 9-labelled XH001/TM7x were then dropped on to the glass bottom dish, gelation happened on top of the coculture aggregates. Imaging before expansion was acquired after gelation step, then water-induced expansion, and imaging after expansion.

### Spectral imaging and linear unmixing

Spectral images were acquired using a Carl Zeiss LSM 780 confocal microscope with a Plan-Apochromat 20×, 0.8 N.A. dry objective or 63×, 1.4 N.A. oil immersion objective. Images were obtained using simultaneous excitation with 405-, 488-, 561-nm laser lines for both dual-species *Streptococcus* biofilms and three-member *F. nucleatum* wildtype*/F. nucleatum* Δ*radD*-*S. mutans*-*S. sanguinis* coaggregation assay. Spectral images were harvested at a bin size of 8.9 nm. Individual reference emission profiles were obtained under the same conditions. Raw spectral images were smoothed by a median filtering algorithm with a kernel pixel size of 3 through ZEN Black software (Carl Zeiss). Linear unmixing was then performed using ZEN Black software (Carl Zeiss) using reference spectra. Unmixed images were assembled and pseudo-colored using Fiji software (52).

## Acknowledgments

This work was supported by National Institutes of Health (NIH) grant T90 DE026110 (to P.-T.D.), Forsyth Pilot Grant FPILOT80 (to P.-T.D.), NIH Grant DE022586 (to J.L.M.W and G.G.B), and NIH Grant DE023810, DE030943 (to X.H). We thank Ms. Jennifer Gundrum for technical assistance, Dr. Jessica L. Mark Welch and Dr. S. Tabita Ramirez-Puebla for helpful discussions.

## Author Contributions

Conceptualization, P.-T.D., and G.G.B.; Investigation, G.G.B., X.H., and W.S.; Writing – Original Draft, P.-T.D., and G.G.B.; Writing – Review & Editing, P.-T.D., W.S., X.H., and G.G.B.; Visualization, P.-T.D.; Supervision, G.G.B., and X.H.; Funding Acquisition, P.-T.D., and G.G.B.

## Supporting information

**Fig. S1.**
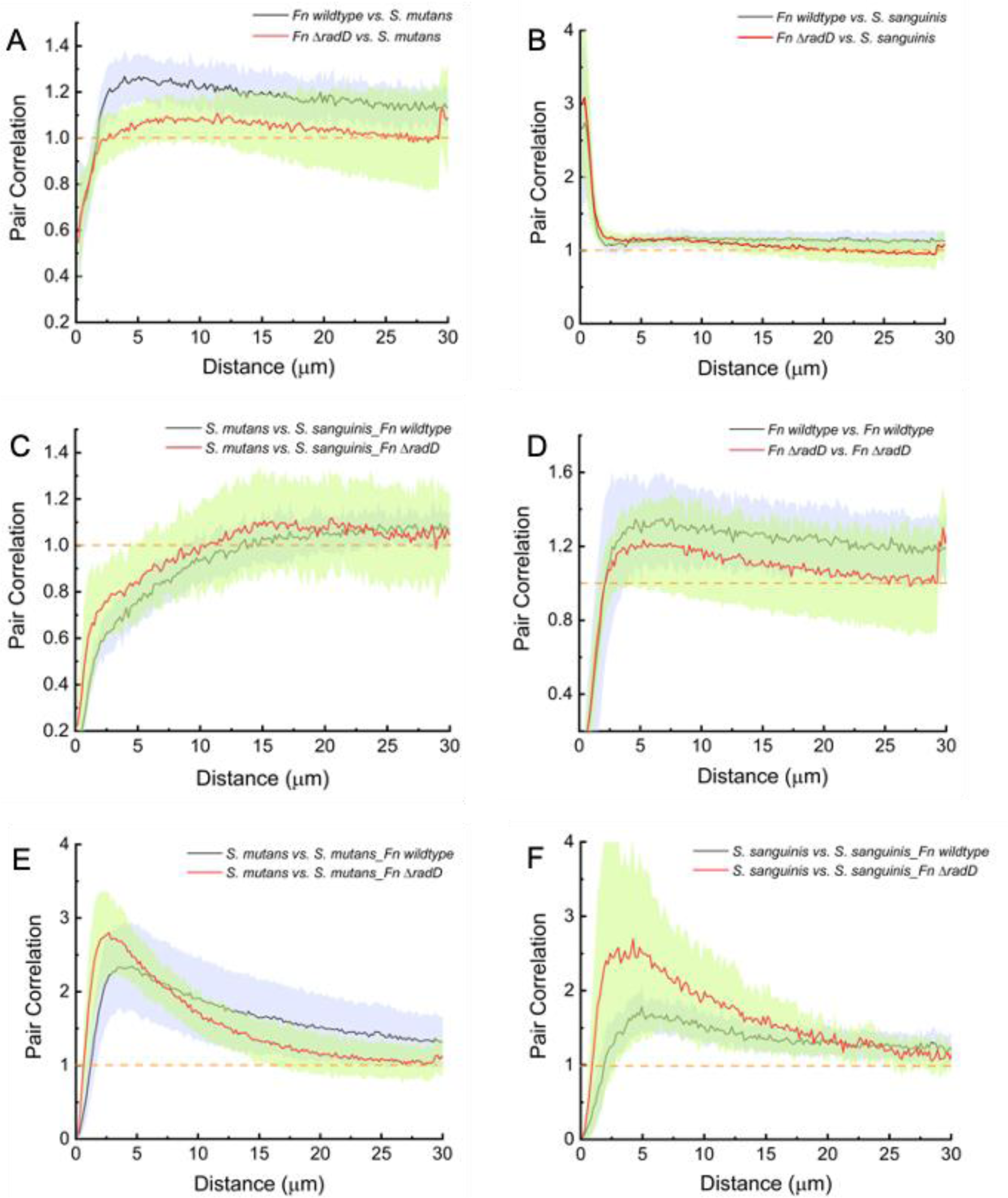
Linear dipole analysis quantifies the spatial organization of taxa within *F. nucleatum*-*S. mutans*-*S. sanguinis* aggregates. Data: Mean (solid line) with 95% confidence interval (shaded area) from 8 different fields of view. Pairwise correlations were calculated using 500,000 random dipoles per field of view. A pair correlation value of 1 is highlighted by orange dashed lines in the figure panels.

**Fig. S2.**
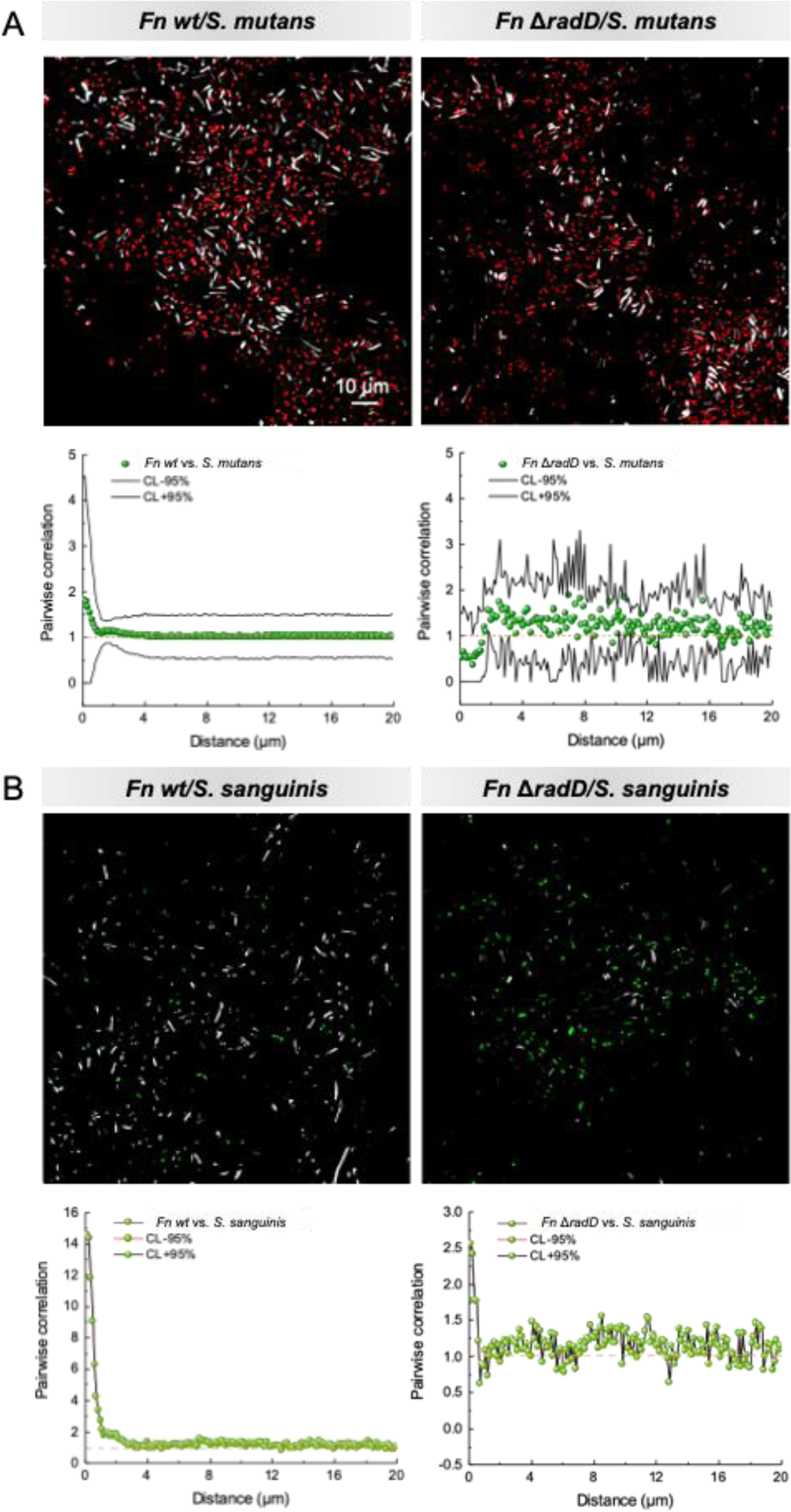
Confocal fluorescence imaging along with corresponding spatial arrangement analysis of aggregates formed between *Fn* and two individual *Streptococcus* species after expansion.

**Fig. S3.**
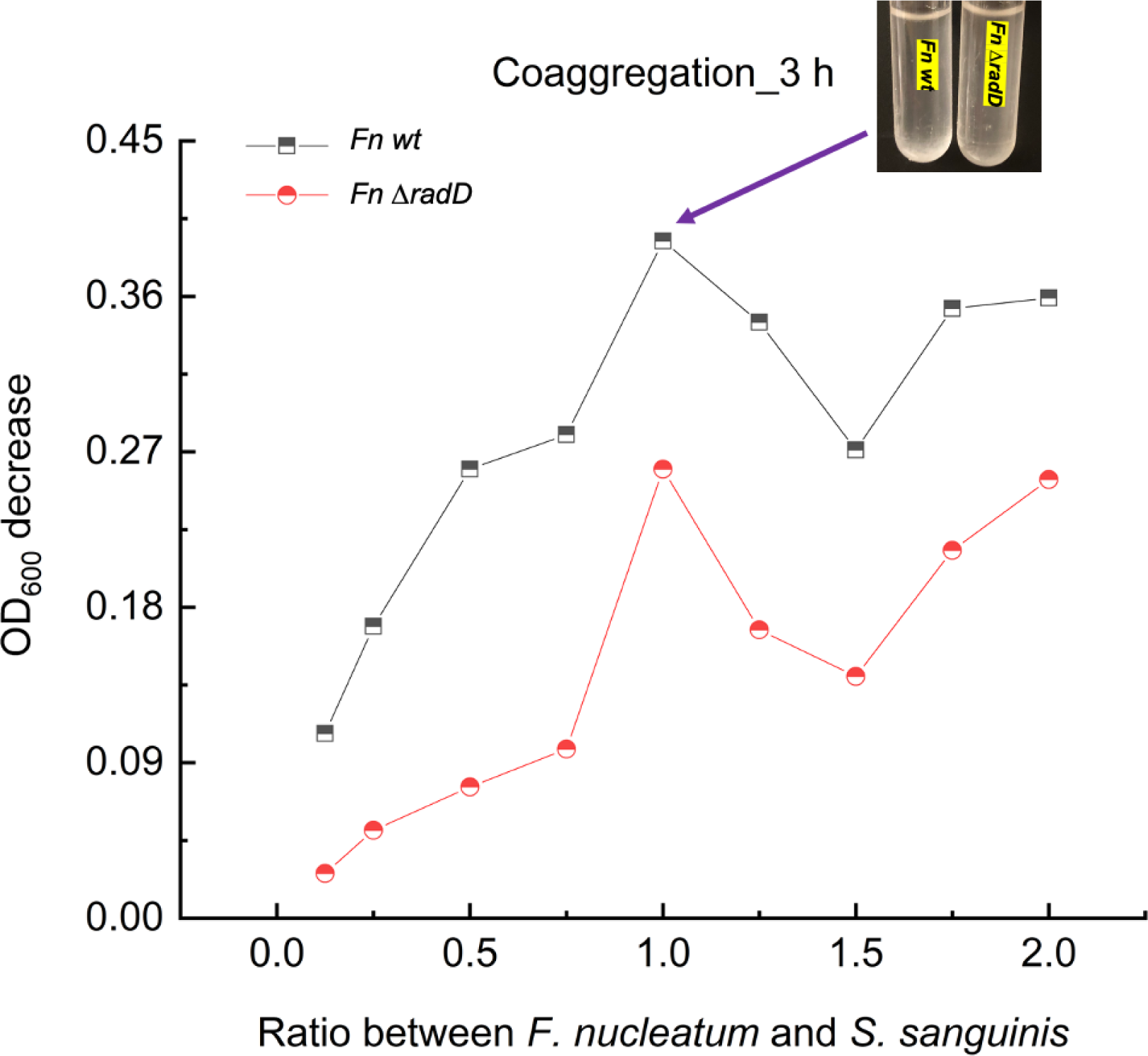
Coaggregation assay between *Fn* wt/*Fn* Δ*radD* and *Streptococcus sanguinis*.

